# Identifying challenges and opportunities for histone peptides in protein engineering platforms

**DOI:** 10.1101/2021.12.28.474372

**Authors:** Jenna N Meanor, Albert J Keung, Balaji M Rao

## Abstract

Histone post-translational modifications are small chemical changes to histone protein structure that have cascading effects on diverse cellular functions. Detecting histone modifications and characterizing their binding partners are critical steps in understanding chromatin biochemistry and have been accessed using common reagents such as antibodies, recombinant assays, and FRET based systems. High throughput platforms could accelerate work in this field, and also could be used to engineer de novo histone affinity reagents; yet published studies on their use with histones have been noticeably sparse. Here we describe specific experimental conditions that affect binding specificities of post-translationally modified histones in classic protein engineering platforms and likely explain the relative difficulty with histone targets in these platforms. We also show that manipulating avidity of binding interactions may improve specificity of binding.

## Introduction

Post-translational modifications (PTMs) of histone proteins play pivotal roles in orchestrating chromatin function including in DNA repair, gene transcription, and cell replication^1–4^. The characterization of natural binders of histone PTMs as well as the generation of engineered affinity reagents such as antibodies has advanced our understanding of chromatin biology^1,5,14,6–13^. However, despite the importance of molecular interactions with histone PTMs, current processes to characterize them and to engineer new affinity reagents are typically laborious, low throughput, and often result in reagents with variable specificity^15^. This is despite the fact that high throughput platforms exist for characterizing and engineering proteins more generally, including platforms such as yeast surface display, phage display, and mRNA display as well as high throughput screening techniques such as magnetic activated cell sorting (MACS) and fluorescent activated cell sorting (FACS)^16,17^.

Here, we identify a critical limitation in the conventional workflow of some of these platform approaches that may explain their underutilization in the context of histone PTMs^18^. The isolation or identification of histone binding proteins using platforms such as yeast surface display typically rely on biotin-mediated immobilization of histone PTM targets on magnetic beads for subsequent panning and magnetic separation of putative binders. Here through a series of experiments we show that immobilization of biotinylated peptides on streptavidin-functionalized magnetic beads results in loss of binding specificity to histone PTMs. We then present an alternative strategy that may alleviate the problems arising from peptide immobilization.

## Results

### Yeast surface display provides a facile platform to characterize histone reader specificities

For this study, we chose six protein domains with a diverse range of specificities and affinities for histone PTMs as reported in the literature: the chromodomain of MPP8, the tandem-Tudor domains (TTDs) of UHRF1, the BAH domain of ASH1L, the bromodomain of ATAD2, the bromodomain of BPTF, and the HDM-JARID domain of KDM5D (**Table 1, Supplemental Table 1)**. We first asked if the binding specificities and relative affinities of these natural binding domains could be readily characterized in a semi-high throughput fashion, without the need for recombinant protein production and purification. We leveraged yeast surface display technology to present the protein domains and mixed the yeast with soluble synthetic peptides with PTMs to quantitatively assess binding specificity (**Supplemental Table 2**). Briefly, yeast cells displaying one of the six proteins were incubated with a titration series of modified histone peptides that were also biotinylated to provide a handle for fluorescent labeling. At each peptide concentration, the fraction of displayed protein bound to the soluble peptide was determined by Streptavidin-PE labeling of the biotinylated peptide through flow cytometry (**Figure 1A, Supplemental Table 3**). Data were fit to a monovalent binding isotherm to estimate apparent equilibrium dissociation constants as previously described^16^ (**Figure 1B-C**). For all proteins, binding data followed expected binding trends based on previous literature (**Table 1**), suggesting this as a facile method for characterizing binding of proteins to histone peptides with PTMs.

**Figure 1.**
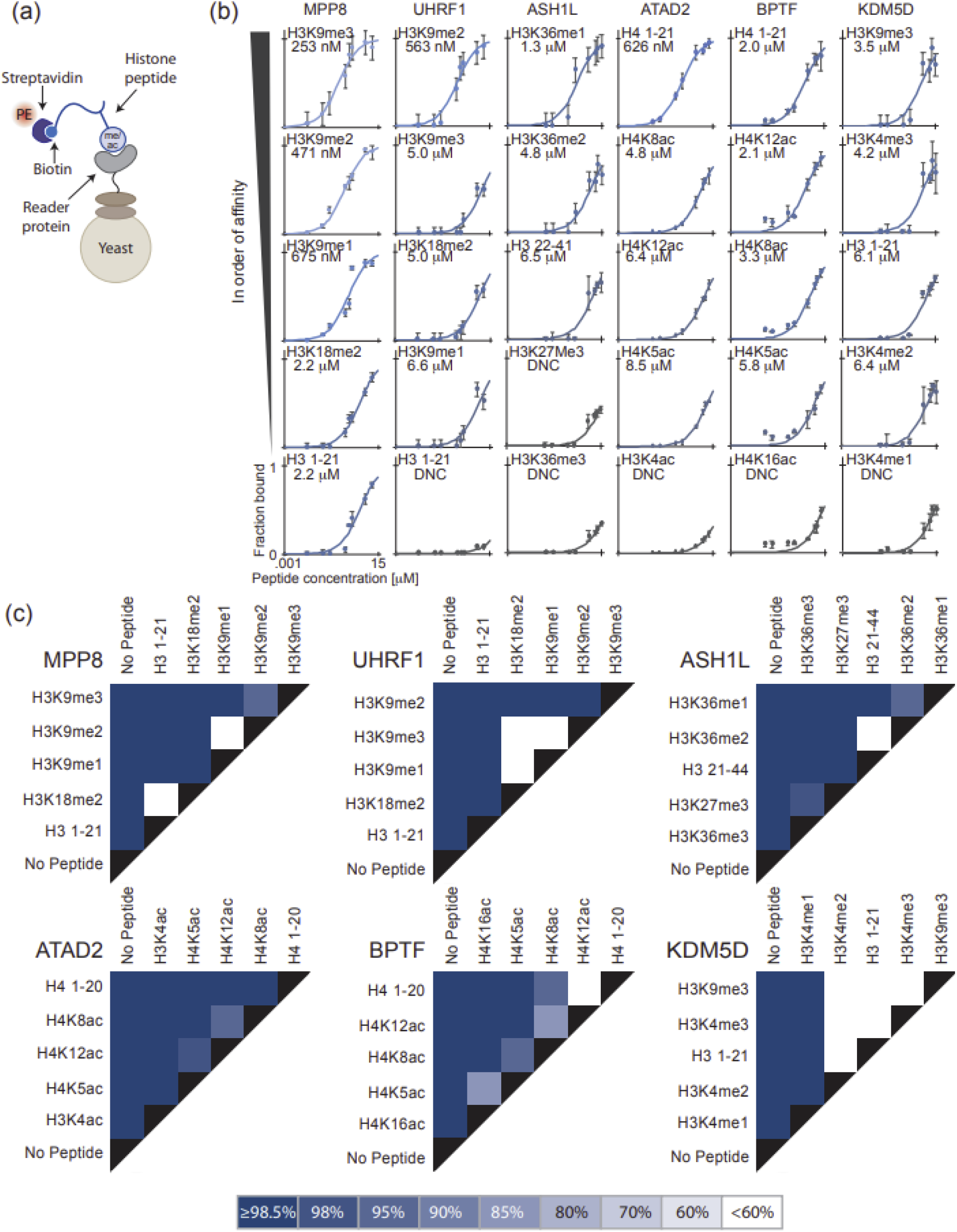
Yeast surface display provides a facile platform to characterize histone reader specificities. (**a**) Interaction between biotinylated peptide and displayed reader protein is measured via streptavidin PE; (**b**) Binding isotherms and calculated binding affinities of displayed reader proteins to respective peptides; error bars represent the standard deviation from triplicate samples; (**c**) Binding affinity discrimination determined by overlap of confidence intervals. More distinct binding affinities (K_Ds_) exhibit higher percentage confidence intervals that do not overlap. Darker colors are associated with a higher level of discrimination between respective peptides. Legend indicates the confidence intervals at which binding affinities were distinguishable.

**Table 1.** Human protein domains used in experiments along with function and histone PTM binding preferences.

| **Protein** | **Domain** | **aa** | **Function** | **Histone PTM binding** |
| --- | --- | --- | --- | --- |
| MPP8<br>19–22 | Chromodomain | 49-120 | Interacts with H3K9 methyltransferases GLP and ESET and DNA methyltransferase 3A | H3K9 methylation |
| UHRF1<br>20,23,24 | Tandem Tudor domains | 127-285 | E3 ubiquitin ligase, recruits DNMT1 | H3K9 methylation |
| ASH1L<br>2,25,26 | BAH | 2261-2798 | H3K36 methyltransferase | H3K36 lower-order methylation |
| ATAD2<br>26–28 | Bromodomain (IV) | 1001-1071 | Interacts with androgen receptor, estrogen receptor alpha, and E2F transcription factors | H4 acetylation |
| BPTF<br>26,28 | Bromodomain (I) | 2944-3014 | Subunit of NURF chromatin-remodeling complex | H4 acetylation |
| KDM5D<br>2,25 | HDM-JARID | 14-55 | H3K demethylase | H3K higher-order methylation |

### Discrimination of binding specificity and detection of weak binders are abrogated in MACS

The individual labeling of yeast displaying histone binding proteins results in specific and clean results and provides a facile and accessible surrogate approach to characterize binding specificities and relative affinities of proteins to histone PTMs. However, scaling this type of characterization to many proteins or panning for specific histone binders from a diverse library of protein candidates requires a different approach; typically, peptides are immobilized on magnetic beads and used to pan binders from a library of candidates (i.e., magnetic activated cell sorting or MACS). Therefore, we next asked if we could identify a potential reason this high throughput approach has not been widely implemented, or at least reported, previously.

To do this, we first tested if the selected protein domains retained PTM-specific binding when the biotinylated histone peptides were immobilized on streptavidin-coated magnetic beads rather than freely presented in solution (**Figure 2A**). These beads were mixed with yeast that both displayed the binder proteins and an engineered luciferase reporter, NanoLuc. The beads and bound yeast were then pulled down using a magnet. The NanoLuc allowed for quantification via luminescence as previously described by Bacon et al^29^. Based on a recent quantitative yeast-yeast two hybrid system, the relative amount of yeast pulled down from the system by each modified histone peptide should rely solely on the strength of the interaction^29^; the stronger the K_D_, the more yeast should be removed. Yeast displaying just luciferase were also tested as a negative control and exhibited similar background to yeast displaying proteins mixed with non-target histone peptides (**Supplemental Figure 1**).

**Figure 2.**
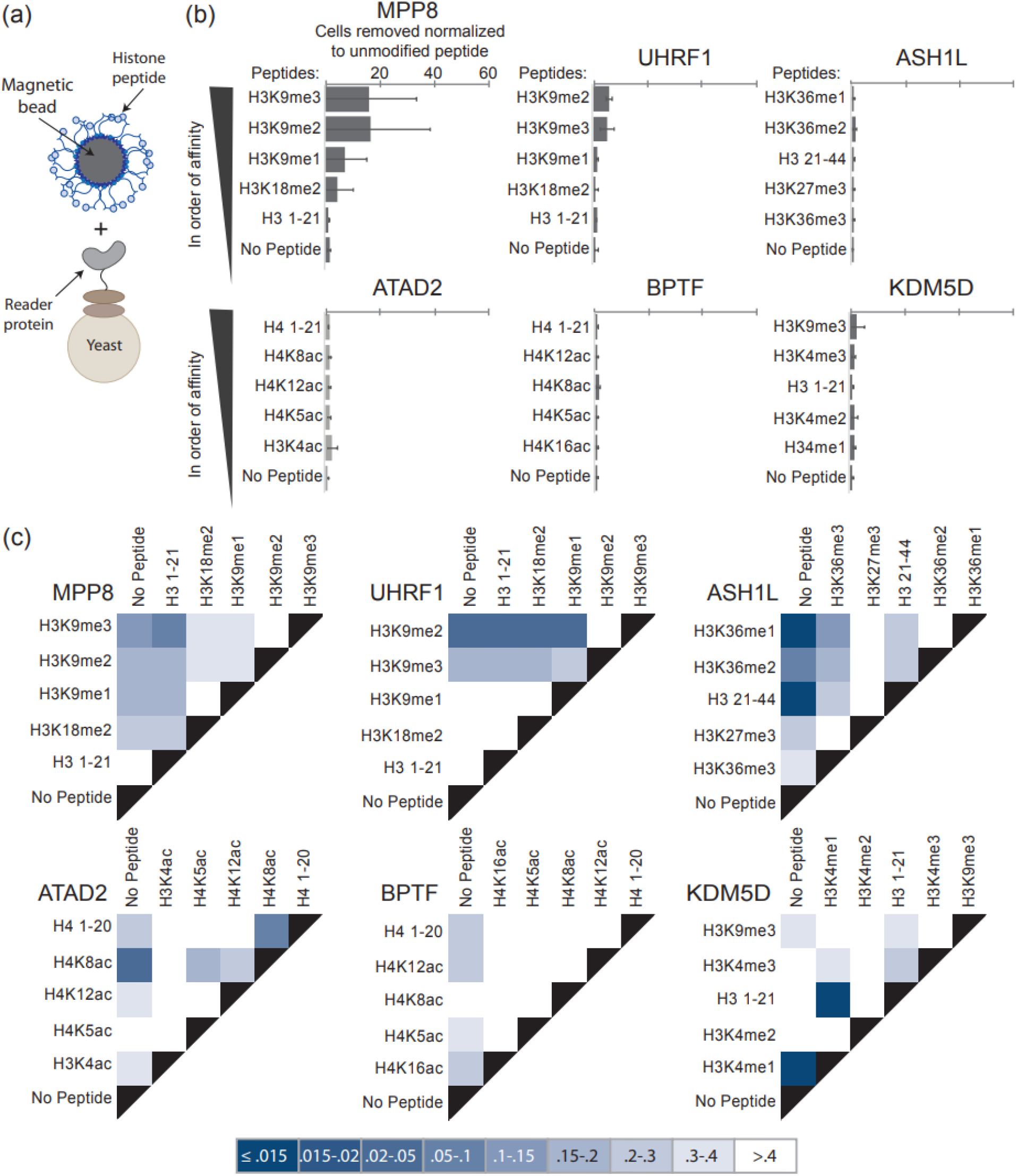
Discrimination of binding specificity and detection of weak binders are abrogated in MACS. (**a**) Interaction between peptide coated magnetic bead and yeast displaying reader protein; (**b**) Relative amount of yeast displaying reader proteins magnetically separated by beads linked to modified histone peptides compared to an unmodified histone peptide control; error bars represent standard deviation from triplicate samples; peptides displayed in order of binding affinities calculated in Figure 1 and from literature; (**c**) Discrimination between amount of yeast separated via magnetization; darker colors are associated with a higher level of discrimination; legend indicates p-value comparing each peptide pair-wise comparison via single factor ANOVA.

Interestingly the number of cells pulled down by each histone peptide, normalized to the number pulled down with an unmodified control histone peptide, did not match expected trends in relative affinities (**Figure 2B, Table 1**). For those proteins with stronger overall affinities to modified histone peptides, such as MPP8 and UHRF1, MACS was unable to distinguish between closely related PTMs. Specifically, for both MPP8 and UHRF1, classic MACS was unable to discriminate binding between H3K9me2 and H3K9me3. Furthermore, there was no discernable pattern for proteins with weaker affinities to their respective PTMs (ASH1L, ATAD2, BRTF, KDM5D), potentially suggesting limitations of this platform to both discriminate binding to specific histone peptides as well as capture low affinity binders in general. The stringency and specificity of pull-down assays are commonly controlled by tuning buffer conditions. We therefore screened a wide range of buffer conditions that varied surfactant and protein concentrations, ionic strength, and yeast to bead ratios. Despite testing many distinct conditions informed by literature, we observed no significant improvement in the binding specificity and ability to capture weak affinity binders (**Supplemental Figure 2**).

### Antibodies label more specifically when histone peptides are presented on yeast versus bead surfaces

We hypothesized that linking peptides to the surface of the beads might be negatively affecting binding specificity. We therefore further challenged peptide-linked beads with a distinct and widely used set of affinity reagents, antibodies, and found they also exhibited poor binding specificity. Streptavidin coated magnetic beads were first linked to biotinylated peptides containing unmodified, mono, di, or tri-methylated lysine 9, and then with corresponding primary and secondary antibodies to each specific modification (**Figure 3A, Supplemental Table 4**) followed by detection by flow cytometry. In all conditions tested, antibodies cross-reacted significantly and non-specifically with all four histone peptides (**Figure 3B, Supplemental Figure 3**).

**Figure 3.**
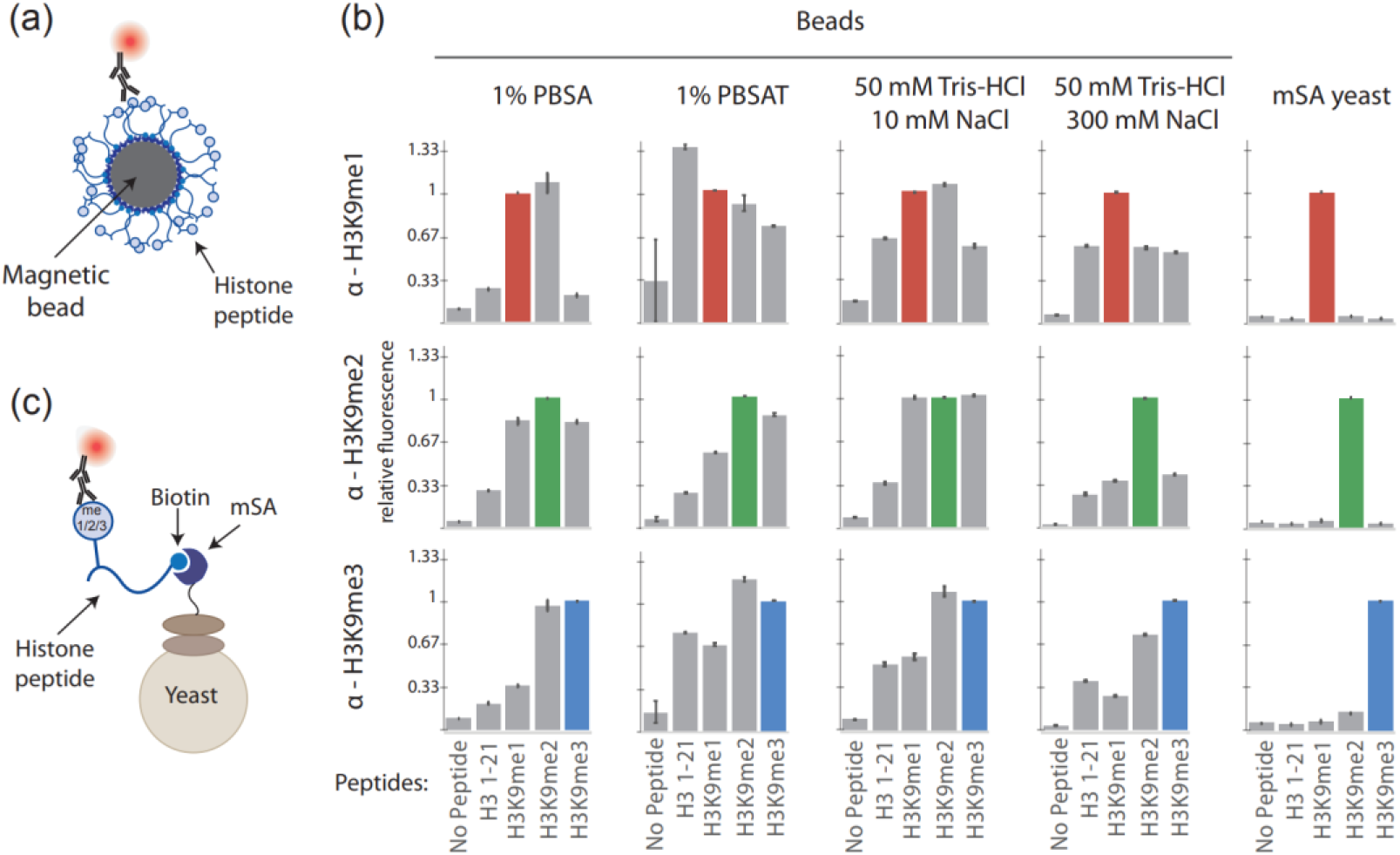
Antibodies label more specifically when histone peptides are presented on yeast versus bead surfaces. (**a**) Interaction between peptide coated magnetic bead and corresponding primary and secondary antibodies; (**b**) Relative fluorescence of each peptide–antibody pairing. The pairings with expected specific interactions with each antibody are indicated by colored bars; error bars represent standard deviation from triplicate samples; (**c**) Interaction between yeast displaying mSA, biotinylated peptide, and corresponding primary and secondary antibodies.

While these antibodies were chosen for their widespread use in many publications, we considered that the antibodies themselves may lack specificity; however, we found that the antibodies were indeed specific when the histone peptides were displayed in a different context. Specifically, when the same set of peptides were linked to the surface of yeast (instead of magnetic beads) through the display of a modified monovalent streptavidin (mSA)^30–32^ (**Figure 3C**), the same antibodies were able to specifically bind to their target epitope and showed significantly less binding to non-target epitopes (**Figure 3B, right**). While the sizes of *Saccharomyces cerevisiae* and the streptavidin coated magnetic beads used in these experiments are on the same order of magnitude, 5 μm diameter and 2.8 μm diameter, respectively, the amount of displayed peptide is not. *S. cerevisiae* can display between 30,000 and 50,000 proteins of interest using the Aga1p and Aga2p display system while magnetic beads can present up to 2 million peptides^16^. This could potentially result in a large avidity effect, masking the ability to distinguish between small differences between histone peptide modifications on beads^33,34^.

### Decreasing peptide density on beads does not rescue antibody labeling specificity

To try and mimic the lower density of peptide achievable on yeast, free biotin and biotinylated H3K9me2 peptide were added in increasing ratios to streptavidin coated magnetic beads while keeping the total biotin content the same; the peptide density on the surface of magnetic beads could be reliably decreased (**Figure 4A**). Two of the lower peptide density conditions were chosen and labeled with antibodies followed by flow cytometry analysis (**Figure 4B**). Even against a significantly decreased surface peptide density on the magnetic beads (16.7 and 3.33%), antibodies were not able to distinguish between unmodified, mono, di, and tri-methylated lysine 9 (**Figure 4C**). Other buffers were also tested but unable to rescue antibody performance (**Supplemental Figure 4**). These results suggest that immobilization density alone cannot fully explain the degraded performance of peptide-labeled beads.

**Figure 4.**
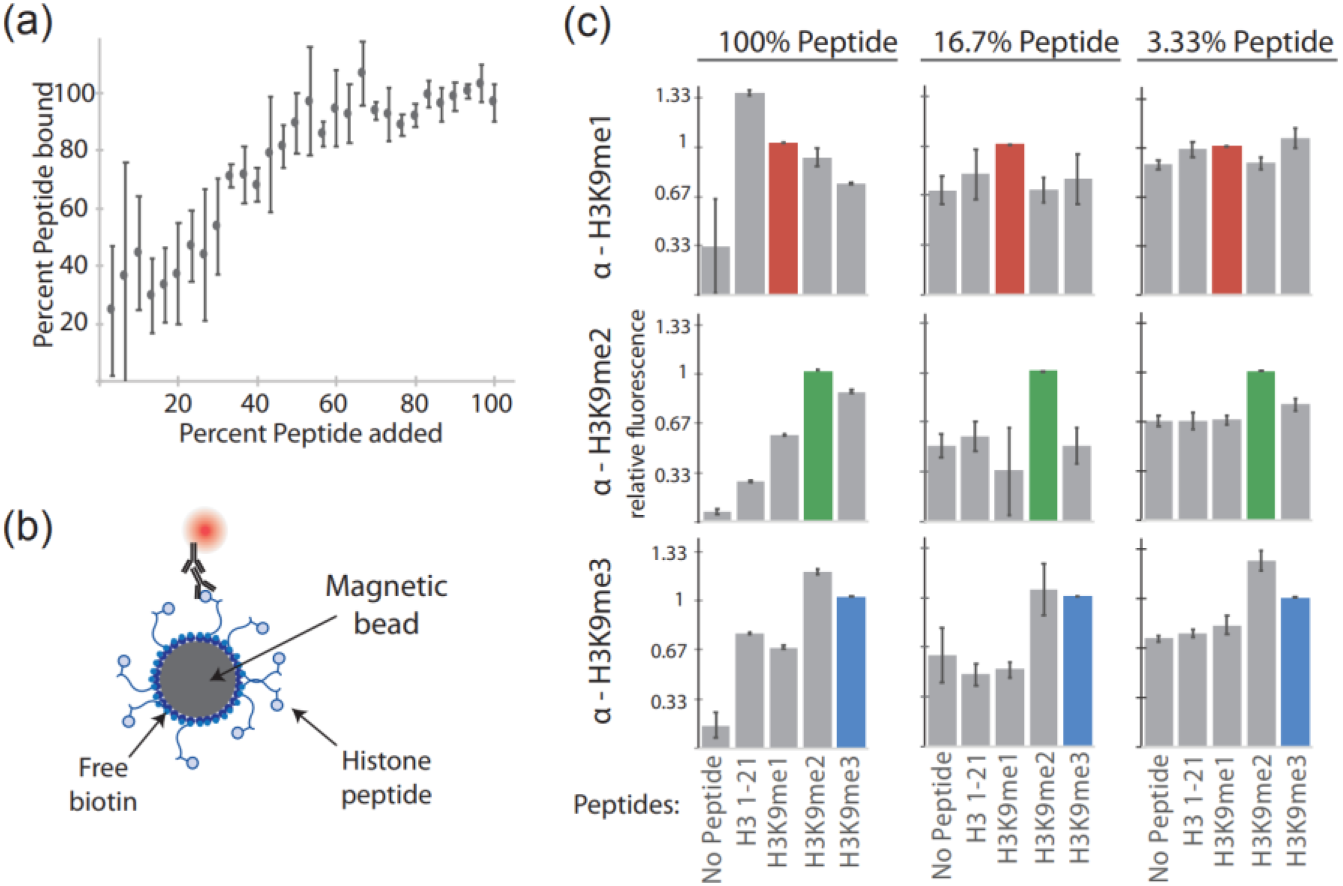
Decreasing peptide density on beads does not rescue antibody labeling specificity. (**a**) Range of H3K9me2 peptide coverage on magnetic beads was achieved by changing free biotin to biotinylated peptide ratio. Percent peptide bound was measured via flow cytometry; error bars represent standard deviation from triplicate samples; (**b**) Interaction between low peptide density magnetic beads and corresponding primary and secondary antibodies; (**c**) Relative fluorescence of each peptide–antibody pairing. The pairings with expected specific interactions with each antibody are indicated by colored bars. Buffer used was 1% PBSAT; error bars represent standard deviation from triplicate samples.

### Soluble peptide binding followed by immobilization improves specificity and yield

As another approach to try and mitigate the negative effects of peptide immobilization on magnetic beads, we tested one more method. This method began with soluble peptide labeling of yeast displaying protein binders (**Figure 5A**). Once the interaction between freely soluble histone peptides and binding proteins displayed on the yeast surface reached equilibrium, excess unbound peptide was washed away. Then, streptavidin coated magnetic beads were introduced to the system. This change in the order of protocol steps (“soluble MACS”) reduces potential unwanted avidity effects in the interaction between displayed protein and biotinylated peptide^35^. For the binding domains that have higher affinities (MPP8 and UHRF1) soluble MACS was able to moderately increase discrimination between modified histone peptides (**Figure 5B-C**). This effect appeared due to a higher yield of cells pulled down in soluble MACS compared to conventional MACS (**Figure 6**). For those domains with relatively weak affinities towards modified histone peptides (ASH1L, ATAD2, BPTF, and KDM5D) even soluble MACS only slightly improved specificity, suggesting a limitation towards detecting weak binders remains.

**Figure 5.**
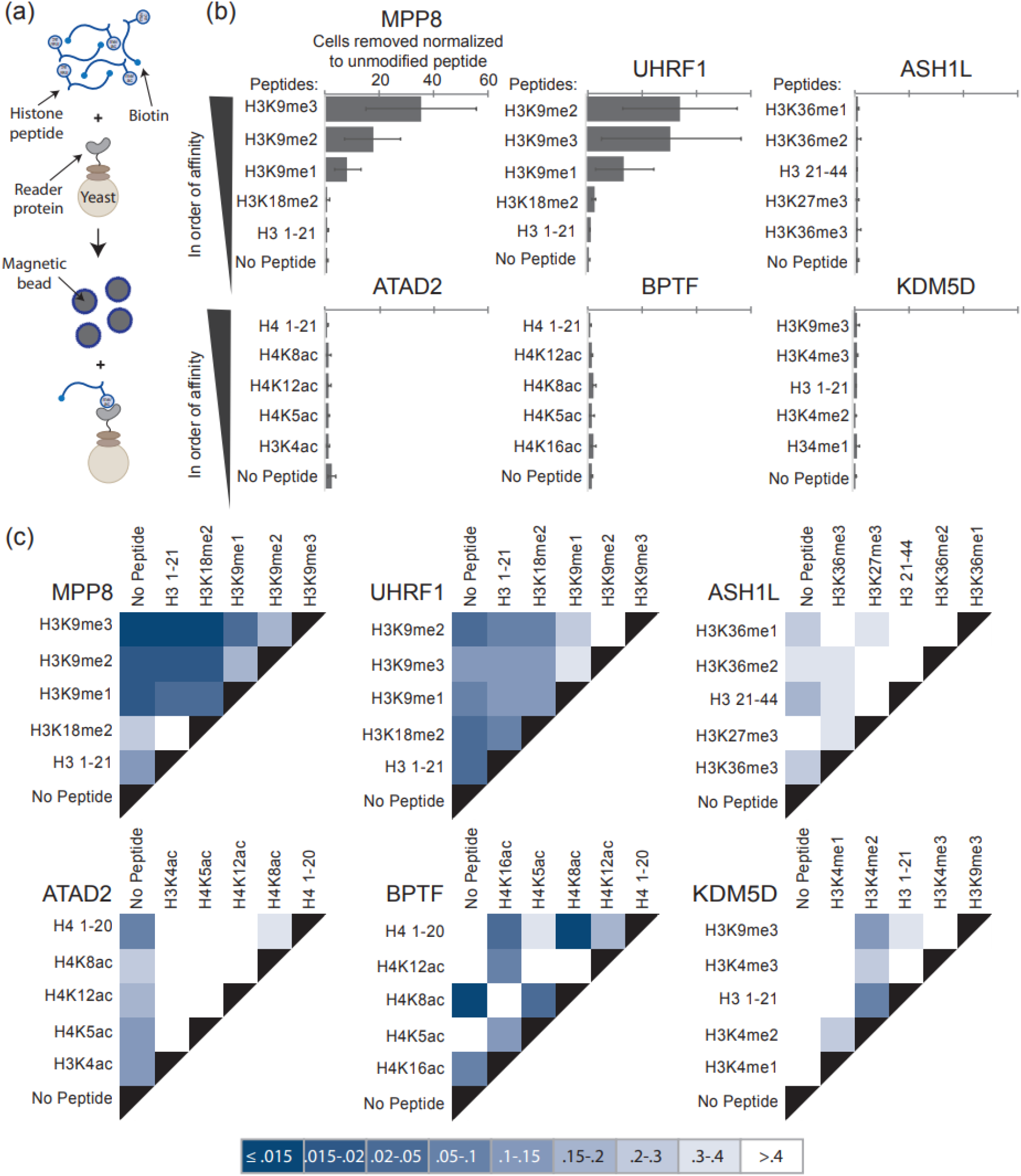
Soluble peptide binding followed by immobilization improves specificity. (**a**) Modified “soluble MACS” method. Interaction between freely soluble biotinylated peptide and yeast displayed protein is then followed by interaction between the peptide-yeast complex to streptavidin coated magnetic beads; (**b**) Relative amount of yeast magnetically separated by modified histones compared to an unmodified peptide control; error bars represent standard deviation from triplicate samples; peptides displayed in order of binding affinities calculated in Figure 1 and from literature; (**c**) Discrimination between amount of yeast separated via magnetization; darker colors are associated with a higher level of discrimination; legend indicates p-value comparing each peptide pair-wise comparison via single factor ANOVA.

**Figure 6.**
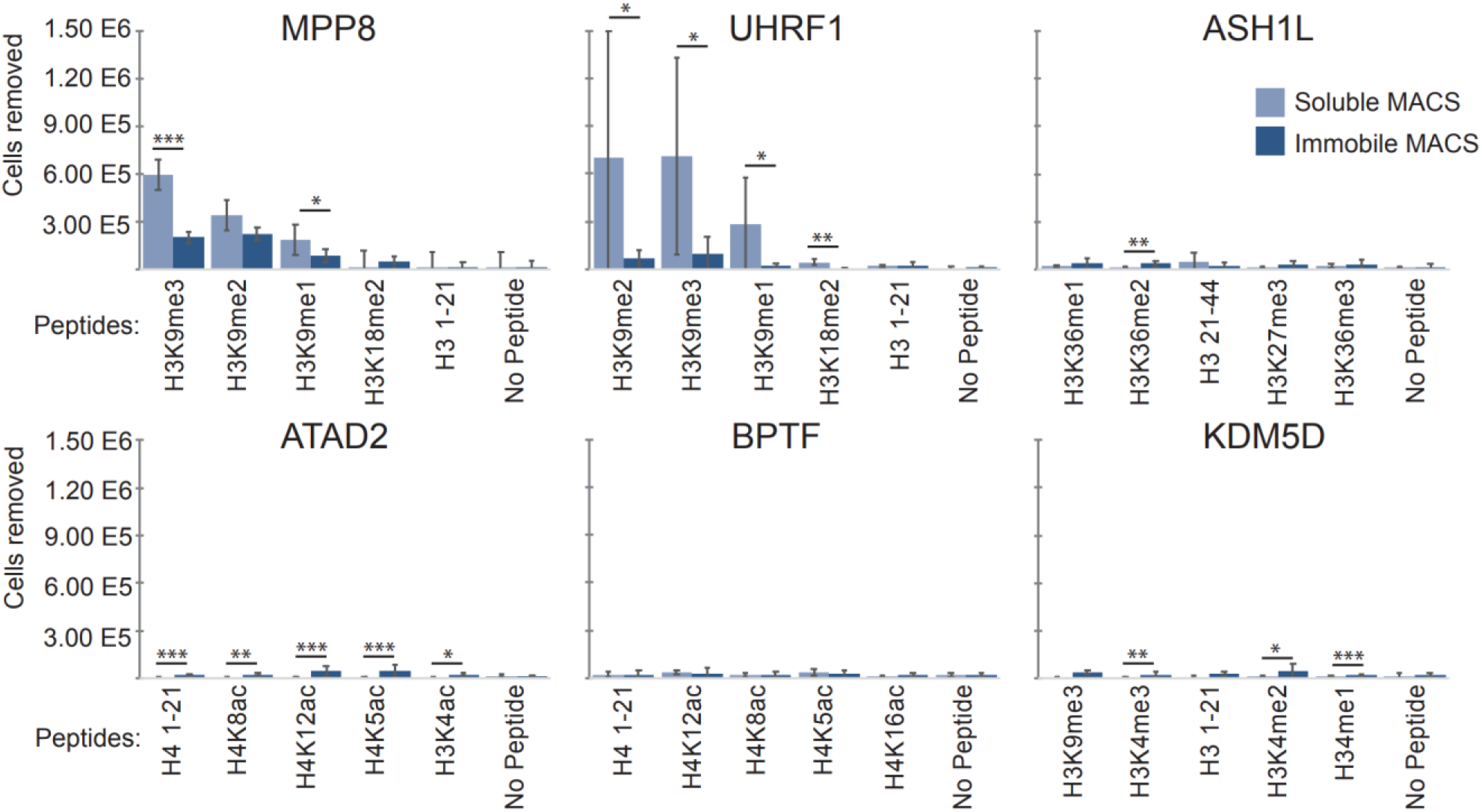
Soluble peptide binding followed by immobilization improves yield. The number of protein displaying cells pulled-down in classic versus soluble MACS methods as a function of histone peptide; error bars represent standard deviation of triplicate pull-downs; \**p*<0.1, \*\**p*<0.05, \*\*\**p*<0.01; *p*-values calculated via single factor ANOVA.

## Discussion

In aggregate, these data suggest that histone peptides do not follow conventional rules when used in traditional protein engineering platforms. Histone tails present several distinct and challenging features. They have high charge density, low hydrophobicity, and are intrinsically disordered^1^. Affinity reagents to histone PTMs must discriminate between differentially modified forms of the same amino acid (H3K9, H3K9me1, H3K9me2, H3K9me3, H3K9ac) where the difference can often be just a few atoms. They would preferably also be able to distinguish the presence or absence of adjacent modifications and distinguish very similar amino acid sequence motifs like H3K9 and H3K27 that both are within an A-R-K-S peptide sequence^1,36^. The mode of attachment of histone peptides to surfaces and their surface density seem to be critical in meeting these challenges. In this work, soluble MACS conditions, which allow for one-to-one interactions between modified histone peptides and proteins displayed on the yeast surface, were more effective in providing this specificity over conventional immobile MACS conditions. However, neither MACS condition could reach the level of specificity achieved by labeling yeast-displayed proteins with soluble histone peptides and analyzing by flow cytometry. Importantly, a broad range of buffer conditions often tuned in other biochemical assays were not able to improve specificity. While flow cytometric approaches can be used in high throughput approaches and directed evolution, future advances that enable the use of MACS with histone targets would unlock the throughput and greater coverage of molecular diversity that MACS affords over fluorescence activated cell sorting. These findings may also have implications in other techniques using immobilized modified histone peptides such as peptide arrays and peptide pull-downs using avidin-agarose beads. The mode of immobilization, immobilization surface properties, and peptide density should be considered as they may have more of an effect on biomolecular recognition specificity than previously considered. These key bottlenecks may explain the dearth of high throughput approaches applied to the characterization or engineering of histone binding proteins and affinity reagents. Future work using biophysical, structural biology, and biochemical characterizations could further elucidate the mechanism(s) for the degraded performance of histone peptides on surfaces and could unlock the full use of high throughput platforms, directed evolution, and combinatorial screening in epigenome engineering.

## Materials and Methods

### Plasmids and yeast culture

The Saccharomyces Cerevisiae strain EBY100 was used for all yeast experiments; pCTCON vector contains a TRP selectable marker and pCT302 vector contains a LEU selectable marker. Plasmid DNA was transformed into chemically competent EBY100 using the Frozen-EZ yeast transformation Kit II (Zymo Research). Trp-deficient SDCAA and SGCAA medium was used for culturing cells and for inducing cell surface protein expression for cells containing the pCTCON vector and Leu-deficient SDSCAA1 (-Leu) and SGSCAA1 (-Leu) media was used for cells containing the pCT302-based vector. Leu- and Trp-deficient SDSCAA2 (-Leu/-Trp) and SGSCAA2 (-Leu/-Trp) media were used for cells containing both the pCT302 and pCTCON-based vectors. (-Leu) and (-Leu/-Trp) media have similar composition to SDCAA and SGCAA media except they contain synthetic dropout mix (1.62 g/L) lacking leucin, or leucine and tryptophan, respectively, instead of casamino acids. Yeast cells cultured in SDCAA, SDSCAA1, or SDSCAA2 medium, as appropriate, for 20-24 hours at 30°C with shaking at 250 rpm. Protein expression was induced by transferring cells into SGCAA, SGSCAA1, or SGSCAA2 medium at an OD_600_ of 1 and cultured for 16-20 hours at 20°C with shaking at 250 rpm. Un-transformed EBY100 was grown in YPD medium for 20-24 hours at 30°C with shaking at 250 rpm.

### Plasmid construction

All displayed proteins were encoded as fusions to Aga2p, a yeast cell mating protein. The chromodomain of MPP8, the tandem-Tudor domains (TTDs) of UHRF1, the BAH domain of ASH1L, the bromodomain of ATAD2, the bromodomain of BPTF, and the HDM-JARID domain of KDM5D were all inserted between the NheI and BamHI sites of pCTCON using amplification primers with restriction enzyme cut sites to generate pCTCON-MPP8, pCTCON-UHRF1, pCTCON-ASH1L, pCTCON-ATAD2, pCTCON-BPTF, and pCTCON-KDM5D. All proteins were amplified from cDNA made from a combination of HEK293T, Jurkat, and K562 cells. Amplified protein and pCTCON backbone were digested with BamHI and NheI restriction enzymes (New England Biolabs) according to the manufacturer’s protocol. Restriction digests were performed in 20 μL for 1 hr at 37°C. Digested plasmid backbones were treated with rSAP (New England Biolabs) for the final 5 minutes of the digestion. Ligations of the digested plasmid backbones and PCR products occurred for 5-10 minutes at RT using T4 DNA ligase (Promega) prior to transformation into NEB^®^ Turbo Competent *E. coli*. 24 hr *E. coli* cultures were harvested for their plasmids using the ZR Plasmid Miniprep – Classic kit. pCT302-NanoLuc plasmid construction is described previously^29^. pYD1-mSA (Addgene plasmid #39865) plasmid construction is described previously^32^.

### Flow cytometry and affinity determination

The equilibrium dissociation constant (KD) was determined using cell surface titrations as described^37^. Briefly, yeast cells displaying one of the histone associating proteins were incubated with various concentration of the biotinylated modified histone peptides in 0.1% PBSA (PBS pH 7.4, 0.1% BSA) at room temperature, followed by Strep-PE. Flow cytometric analysis was used to measure the PE fluorescence intensity for each peptide concentration. The binding affinity between each protein-peptide pairing and confidence intervals were estimated as previously described^16^.

### Magnetic activated cell sorting and luciferase quantification

Magnetic beads were functionalized with biotinylated modified histone peptides by incubating 2 μg of biotinylated peptide per 25 μL Dynabeads Biotin Binder Beads (4 x 10^8^ beads/mL, Thermo-fisher Scientific) for 2 hours with rotation at RT in 0.1% PBSA. Next, the magnetic beads were washed two times with 0.1% PBSA and blocked in 1% PBSA (PBS pH 7.4, 1% BSA) for one hour with rotation at RT. Beads not functionalized with peptide were similarly washed and blocked. 5 x 10^6^ beads were then incubated with 5 x 10^6^ protein and NanoLuc displaying yeast and 5 x 10^8^ EBY100 cells in 2 mL 1% PBSAT (PBS pH 7.4, 1% BSA, 0.05% Tween-20) for 2 hours with rotation at RT. After, the incubations were placed onto a magnet to isolate any cells bound to the magnetic beads. Other incubations buffers tested along with 1% PBSAT were 1% PBSA and heparin buffer (20 mM HEPES pH 7.4, 150 mM KCl, 2 mM MgCl_2_, 2 μg/mL heparin). A one-to-one ratio of beads to displaying yeast is described above and five-to-one and ten-to-one ratios were also tested. EBY100 was always present in 100-fold excess of displaying yeast in all experimental variations.

After any cells not bound to the magnetic beads were removed with a magnet, the beads and bound protein displaying cells were washed gently three times with 1% PBSAT, or respective buffer, and then resuspended in 100 μL of PBS. 100 μL of the Nano-Glo Luciferase Assay system (Promega) was added to the magnetic bead solution. The reaction was allowed to proceed for three minutes and then the tube containing the magnetic beads was placed on to a magnet. 100 μL of the reaction was plated in duplicate onto a 96 black-well plate with a clear, flat bottom. The luminescence was read using a Tecan Infinite 200 plate reader using an integration time of 400 ms, settle time of 0 ms, and no attenuation. Standard curves generated using known quantities of protein displaying cells were used to estimate the number of cells removed with the magnet. p-values were calculated using single factor ANOVA in excel.

### Antibody specificity through flow cytometry on magnetic beads

Magnetic beads were functionalized with biotinylated modified histone peptides as described above. 5 x 10^6^ beads were then incubated with either a H3K9me1, H3K9me2, or H3K9me3 antibody for 30 minutes with rotation at 4°C in 100 μL of incubation buffer (ab1220 1:200, ab9045 1:200, ab8898 1:250, Abcam). Incubation buffers tested were 0.1% PBSA, 1% PBSA, 1% PBSAT, 50 mM Tris-HCl + 10 mM NaCl, 50 mM Tris-HCl + 300 mM NaCl, and 50 mM Tris-HCl + 500 mM NaCl. Samples were then washed with incubation buffer and incubated with secondary antibody for 10 minutes with rotation at 4°C in the dark in 100μL 0.1% PBSA (ab150075 1:250, ab150107 1:250, Abcam). Samples were washed in 0.1% PBSA and run on MACSQuant VYB cytometer using the 561 nm laser and 661/20nm filter. Flow cytometry data was analyzed with FlowJo software.

### Antibody specificity through flow cytometry on mSA displaying yeast

2 x 10^6^ pYD1-mSA containing yeast cells were washed and pelleted prior to labeling. Samples were then incubated with either a H3K9me1, H3K9me2, or H3K9me3 antibody for 30 minutes with rotation at 4°C in 100 μL of 0.1% PBSA (ab1220 1:200, ab9045 1:200, ab8898 1:250, Abcam). Samples were then washed with 0.1% PBSA and incubated with secondary antibody for 10 minutes with rotation at 4°C in the dark in 100μL 0.1% PBSA (ab150075 1:250, ab150107 1:250, Abcam). Samples were washed in 0.1% PBSA and run on MACSQuant VYB cytometer using the 561 nm laser and 661/20nm filter. Flow cytometry data was analyzed with FlowJo software.

### Decreasing peptide density on magnetic beads

Magnetic beads were functionalized with mixtures of biotinylated modified histone peptides and biotin. The total mass of biotin and biotinylated modified histone peptide was kept constant at 1.5 μg. 30 different ratios were tested. For each ratio, 5 x 10^6^ beads were then incubated with an H3K9me2 antibody for 30 minutes with rotation at 4°C in 100 μL of 1% PBSA (ab1220 1:200, Abcam). Samples were then washed with 1% PBSA and incubated with secondary antibody for 10 minutes with rotation at 4°C in the dark in 100μL 0.1% PBSA (ab150107 1:250, Abcam). Samples were washed in 0.1% PBSA and run on MACSQuant VYB cytometer using the 561 nm laser and 661/20nm filter. Flow cytometry data was analyzed with FlowJo software.

Antibody specificity flow as described above was repeated for samples containing 16.7% peptide in both 1% PBSA and 50 mM Tris-HCl + 300 mM NaCl and 3.33% peptide in 1% PBSA.

### Soluble magnetic activated cell sorting and luciferase quantification

1 μg of biotinylated modified histone peptide was incubated with 5 x 10^6^ protein and NanoLuc displaying yeast and 5 x 10^8^ EBY100 cells in 2 mL 1% PBSAT (PBS pH 7.4, 1% BSA, 0.05% Tween-20) for 2 hours with rotation at RT. Tubes were pelleted and washed to remove any unbound peptide and resuspended in 2 mL 1% PBSAT. 5 x 10^6^ washed magnetic beads were added and incubated with cells for 10 minutes with rotation at RT. After, the incubations were placed onto a magnet to isolate any cells bound to the magnetic beads. Luminescence determination proceeded as previously described.

## Supporting Information

Table S1: DNA sequences of protein domains; Table S2: Histone peptides; Table S3: Fluorophore conjugated secondary antibodies and streptavidin; Table S4: Primary antibodies; Figure S1: MACS with yeast displaying NanoLuc; Figure S2: MACS with yeast displaying MPP8 under varying buffer conditions and yeast-to-bead ratios; Figure S3: Antibody labeling of histone peptides presented on magnetic beads; Figure S4: Antibody labeling of low density histone peptides presented on magnetic beads.

## Acknowledgements

This work was supported by the NSF Emerging Frontiers in Research and Innovation program (EFMA1830910), the National Institute of Biomedical Imaging and Bioengineering and the National Institute On Drug Abuse of the National Institutes of Health under Award Numbers R21EB023377 and DP1DA044359, respectively. The content is solely the responsibility of the authors and does not necessarily represent the official views of the sponsors.

