## Supplemental Tables 1-4, Supplemental Figures 1-4 for "Identifying challenges and opportunities for histone peptides in protein engineering platforms"

**Contents:**

Supplemental Table 1: DNA sequences of protein domains

Supplemental Table 2: Histone peptides

Supplemental Table 3: Fluorophore conjugated secondary antibodies and streptavidin

Supplemental Table 4: Primary antibodies

Supplemental figure 1 MACS with yeast displaying NanoLuc

Supplemental figure 2 MACS with yeast displaying MPP8 under varying buffer conditions and yeast-to-bead ratios

Supplemental Figure 3 Antibody labeling of histone peptides presented on magnetic beads

Supplemental Figure 4 Antibody labeling of low density histone peptides presented on magnetic beads

**Supplemental Table 1** DNA sequences of protein domains

| Protein | Domain | DNA Sequence |
|---|---|---|
| MPP8 | Chromodomain | GGCGACAGTGAGGAGGACGGAGAGGATGTGTTCGAGGTGGAGAAGATCC<br>TGGACATGAAGACCGAGGGGGGTAAAGTTCTTTACAAAGTTCGCTGGAAA<br>GGCTATACATCGGATGATGATACCTGGGAGCCCGAGATTCACCTGGAGGA<br>CTGTAAAGAAGTGCTTCTTGAATTTAGGAAGAAAATTGCAGAGAACAAAGC<br>CAAAGCAGTCAGGAAG |
| UHRF1 | Tandem Tudor domains | GACATGTGGGATGAGACGGAATTGGGGCTGTACAAGGTCAATGAGTACGT<br>CGATGCTCGGGACACGAACATGGGGGCGTGGTTTGAGGCGCAGGTGGTC<br>AGGGTGACGCGGAAGGCCCCCTCCCGGGACGAGCCCTGCAGCTCCACGT<br>CCAGGCCGGCGCTGGAGGAGGACGTCATTTACCACGTGAAATACGACGA<br>CTACCCGGAGAACGGCGTGGTCCAGATGAACTCCAGGGACGTCCGAGCG<br>CGCGCCCGCACCATCATCAAGTGGCAGGACCTGGAGGTGGGCCAGGTGG<br>TCATGCTCAACTACAACCCCGACAACCCCAAGGAGCGGGGCTTCTGGTAC<br>GACGCGGAGATCTCCAGGAAGCGCGAGACCAGGACGGCGCGGGAACTCT<br>ACGCCAACGTGGTGCTGGGGGATGATTCTCTGAACGACTGTCGGATCATC<br>TTCGTGGACGAAGTCTTCAAGATTGAGCGG |
| ASH1L | BAH | TTGCTGCTTCGTCAGGGTGACTGTGTGTATCTGATGAGGGATAGTCGGCG<br>CACCCCTGATGGCCACCCGGTCCGTCAGTCCTATCGACTGTTATCTCACAT<br>TAACCGAGATAAACTTGACATCTTTCGCATTGAGAAGCTTTGGAAGAATGAA<br>AAAGAGGAACGGTTTGCCTTTGGTCACCATTATTTCCGTCCCCACGAAACA<br>CACCACTCTCCATCCCGTCGGTTCTATCATAATGAACTATTTCGGGTGCCA<br>CTCTATGAGATCATTCCCTTGGAGGCTGTAGTGGGGACCTGCTGTGTGTTG<br>GACCTTTATACGTATTGTAAAGGGAGACCCAAAGGAGTAAAGGAGCAAGAT<br>GTGTACATCTGTGATTATCGGCTTGACAAGTCAGCACACCTGTTTTACAAGA<br>TCCAC |
| ATAD2 | Bromodomain (IV) | GCTATTGACAAGCGATTCCGAGTGTTTACTAAGCCTGTTGACCCTGATGAG<br>GTTCCTGATTATGTCACTGTAATAAAGCAACCAATGGACCTTTCATCTGTAA<br>TCAGTAAAATTGATCTACACAAGTATCTGACTGTGAAAGACTATTTGAGAGA<br>TATTGATCTAATCTGTAGTAATGCCTTAGAATACAATCCAGATAGAGATCCT<br>GGAGAT |
| BPTF | Bromodomain (I) | CAGGCCCATAAGATGGCCTGGCCTTTCCTTGAACCAGTAGACCCTAATGAT<br>GCACCAGATTATTATGGTGTTATTAAGGAACCTATGGACCTTGCCACCATG<br>GAAGAAAGAGTACAAAGACGATATTATGAAAAGCTGACGGAATTTGTGGCA<br>GATATGACCAAAATTTTTGATAACTGTCGTTACTACAATCCAAGTGACTCCC<br>CATTTTAC |
| KDM5D | HDM-JARID | TGCCCGGTTTTTGAGCCTAGCTGGGCTGAATTCCAAGACCCGCTTGGCTAC<br>ATTGCGAAAATAAGGCCCATAGCAGAGAAGTCTGGCATCTGCAAAATCCGC<br>CCACCCGCGGATTGGCAGCCTCCT |

**Supplemental Table 2** Histone peptides

| Peptide | Sequence | Manufacturer | Catalog # |
|---|---|---|---|
| H3 1-21 | ARTKQTARKSTGGKAPRKQLA - GG - K(BIOTIN) - NH2 | Anaspec | AS-61702 |
| H3 21-44 | ATKAARKSAPATGGVKKPHRYRPG-GK(Biotin) | Anaspec | AS - 64440-025 |
| H3K4Ac | ART-Kac-QTARKSTGGKAPRKQLA - GGYK(Biotin) - NH2 (1-21) | Active motif | 81039 |
| H3K4me1 | ART-K(me1)-QTARKSTGGKAPRKQLA-GGK(Biotin) | Anaspec | AS - 64355 - 025 |
| H3K4me2 | ART - K(me2) - QTARKSTGGKAPRKQLA - GGK(Biotin) | Anaspec | AS-64356-025 |
| H3K4me3 | ART-K(me3)-QTARKSTGGKAPRKQLA-GGK(Biotin) | Anaspec | AS - 64357 - 025 |
| H3K9me1 | ARTKQTAR - K(me1) - STGGKAPRKQLA - GGK(Biotin) | Anaspec | AS-64358-025 |
| H3K9me2 | ARTKQTAR - K(me2) - STGGKAPRKQLA - GGK(Biotin) | Anaspec | AS-64359-1 |
| H3K9me3 | ARTKQTAR - K(me3) - STGGKAPRKQLA - GGK(Biotin) | Anaspec | AS-64360-025 |
| H3K18me2 | ARTKQTARKSTGGKAPR - K(me2) - QLA - GGK(Biotin) - NH2 | Anaspec | AS-64620-025 |
| H3K27me3 | ATKAAR-K(Me3)-SAPATGGVKKPHRYRPG-GK(Biotin) | Anaspec | AS - 64367-025 |
| H3K36me1 | ATKAARKSAPATGGV - K(me1) - KPHRYRPG - GK(Biotin) | Anaspec | AS - 64368 - 025 |
| H3K36me2 | RKSAPATGGV - K(me2) - KPHRYRPGTV - K(BIOTIN) | Anaspec | AS-64601-025 |
| H3K36me3 | ATKAARKSAPATGGV-K(me3)-KPHRYRPG-GK(Biotin) | Anaspec | AS - 64441 - 025 |
| H4 1-20 | SGRGKGGKGLGKGGAKRHRKVLRGG-YK(Biotin)-NH2 | Active motif | 81109 |
| H4K5ac | SGRG - K(Ac) - GGKGLGKGGAKRHRKVLRDNGSGS - K(Biotin) | Anaspec | AS-65229-1 |
| H4K8ac | SGRGKGG - K(Ac) - GLGKGGAKRHRKVLRDNGSGS - K(Biotin) | Anaspec | AS-65230-1 |
| H4K12ac | SGRGKGGKGLG - K(Ac) - GGAKRHRKVLRDNGSGS - K(Biotin) | Anaspec | AS-65208-1 |
| H4K16ac | SGRGKGGKGLGKGGA - K(Ac) - RHRKVLRDNGSGS - K(Biotin) | Anaspec | AS-65209-1 |

**Supplemental Table 3** Fluorophore conjugated secondary antibodies and streptavidin

| Secondary Antibody | Manufacturer | Catalog # | Dilution Used |
|---|---|---|---|
| Goat anti-Chicken IgY (H&L) DyLight®488 Conjugate | Immunoreagents | GtxCk-003-D488NHSX | 1:250 |
| Donkey anti-Rabbit IgG (H&L) DyLight®633 Conjugate | Immunoreagents | DkxRb-003-E633NHSX | 1:250 |
| Donkey anti-Rabbit IgG (H&L) AlexaFluor®647 Conjugate | Abcam | ab150075 | 1:250 |
| Donkey anti-Mouse IgG (H&L) AlexaFluor®647 Conjugate | Abcam | ab150107 | 1:250 |
| Streptavidin, R-Phycoerythrin Conjugate (SAPE) | ThermoFisher | S866 | 1:250 |

**Supplemental Table 4** Primary antibodies

| Antibody Target | Manufacturer | Catalog # | Host Species | Dilution Used |
|---|---|---|---|---|
| HA | Thermo Fischer Scientific | PA1-985 | Rabbit | 1:100 |
| C-Myc | Thermo Fischer Scientific | A-21181 | Chicken | 1:100 |
| H3K9me1 | Abcam | ab9045 | Rabbit | 1:200 |
| H3K9me2 | Abcam | ab1220 | Mouse | 1:200 |
| H3K9me3 | Abcam | ab8898 | Rabbit | 1:200 |

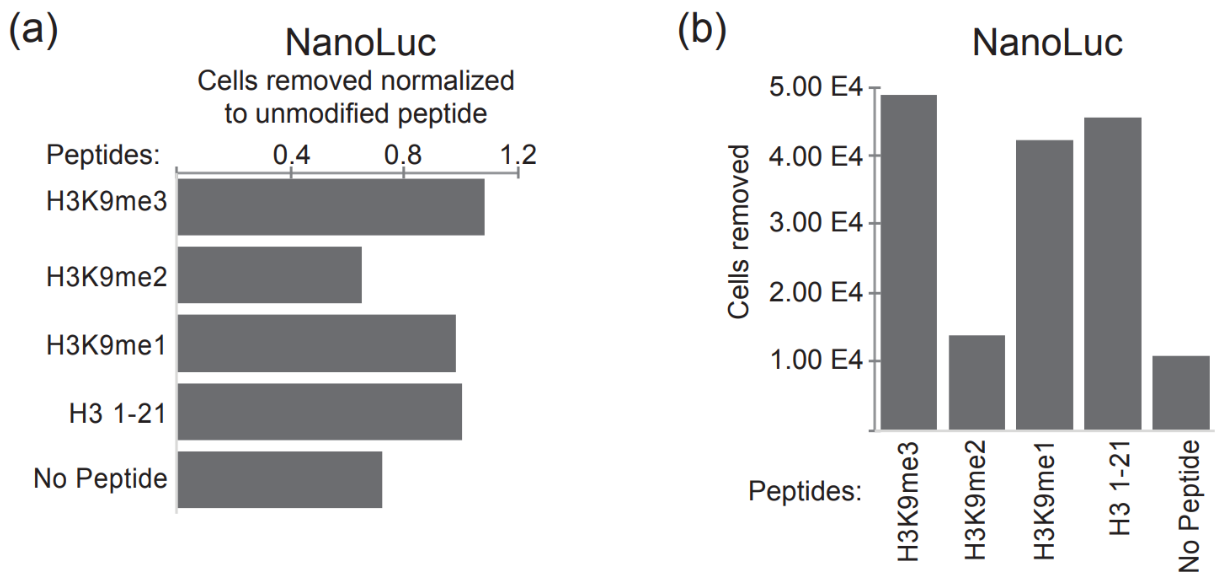

**Supplemental Figure 1.** MACS with yeast displaying NanoLuc (**a**) Relative amount of yeast displaying NanoLuc magnetically separated by beads linked to modified histone peptides compared to an unmodified histone peptide control; (**b**) The number of yeast displaying NanoLuc pulled-down in classic MACS as a function of histone peptide

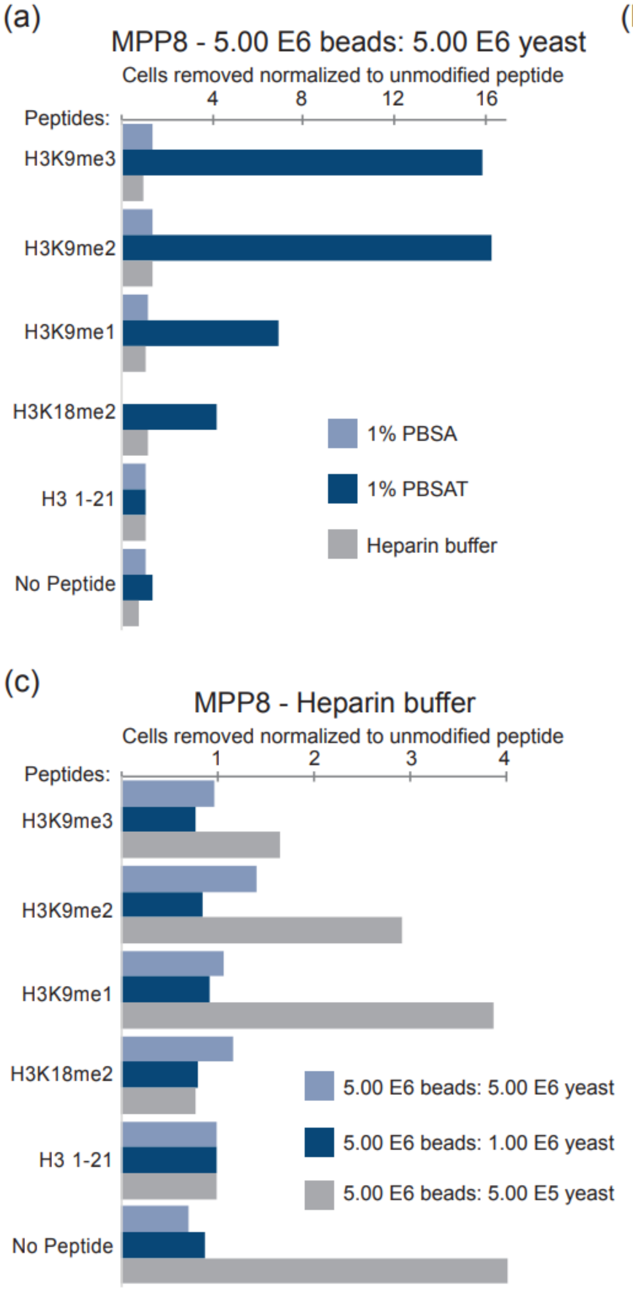

**Supplemental Figure 2** MACS with yeast displaying MPP8 under varying buffer conditions and yeast-to-bead ratios (**a**) Relative amount yeast displaying MPP8 magnetically separated by histone peptide linked to magnetic bead compared to an unmodified peptide control for 1% PBSA, 1% PBSAT, and heparin buffer at a yeast to beads ratio of one-to-one; (**b**) Relative amount of MPP8 displaying yeast magnetically separated by histone peptide linked to magnetic bead compared to an unmodified peptide control in 1% PBSAT for yeast to beads ratios of one-to-one and one-to-five **c** Relative amount of MPP8 displaying yeast magnetically separated by histone peptide linked to magnetic bead compared to an unmodified peptide control in heparin buffer for yeast to beads ratios of one-to-one and one-to-five, and one-to-ten.

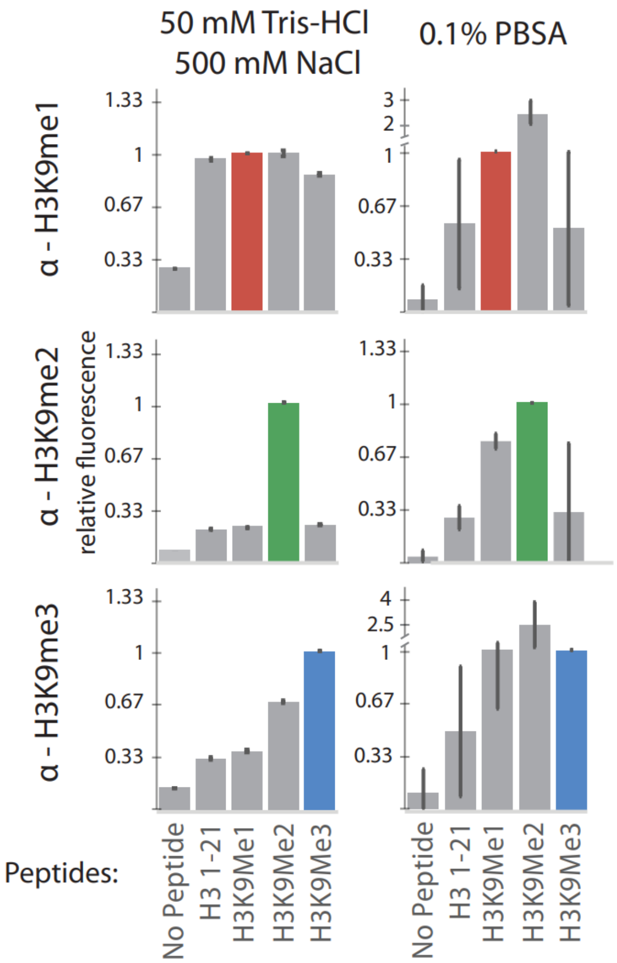

**Supplemental Figure 3** Antibody labeling of histone peptides presented on magnetic beads. Relative fluorescence of each peptide–antibody pairing. The pairings with expected specific interactions with each antibody are indicated by colored bars; error bars represent standard deviation from triplicate samples.

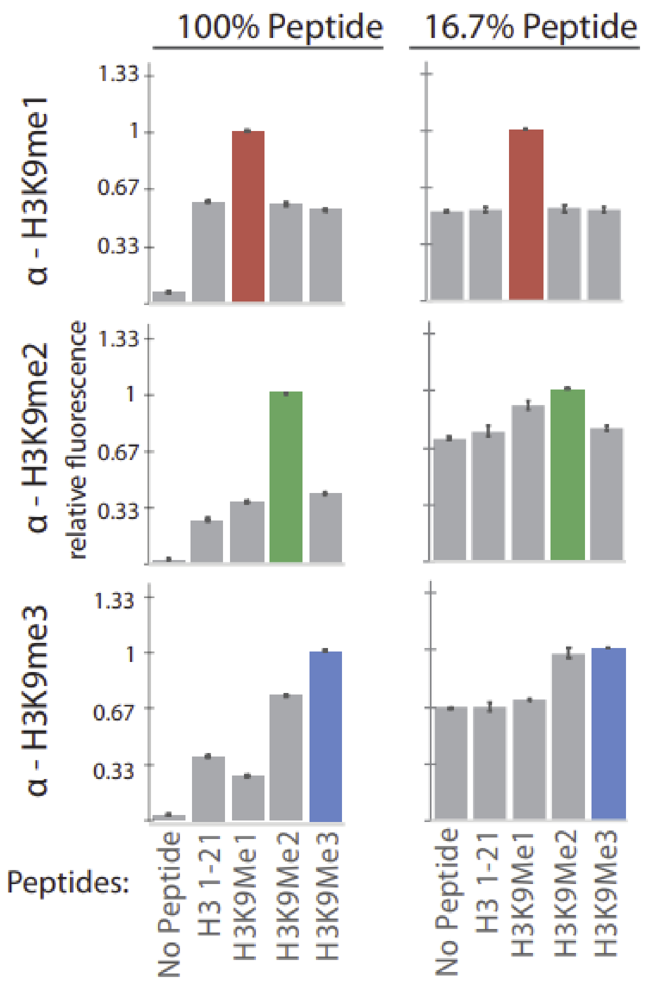

**Supplemental Figure 4** Antibody labeling of low density histone peptides presented on magnetic beads. Relative fluorescence of each peptide–antibody pairing. The pairings with expected specific interactions with each antibody are indicated by colored bars. Buffer used was 50 mM Tris HCl 300 mM NaCl; error bars represent standard deviation from triplicate samples.
